# The synaptic basis of activity-dependent eye-specific competition

**DOI:** 10.1101/2021.09.18.460903

**Authors:** Chenghang Zhang, Colenso M. Speer

## Abstract

Binocular vision requires proper developmental wiring of eye-specific inputs to the brain. Axons from the two eyes initially overlap in the dorsal lateral geniculate nucleus and undergo activity-dependent competition to segregate into target domains. The synaptic basis of such refinement is unknown. Here we used volumetric super-resolution imaging to measure the nanoscale molecular reorganization of developing retinogeniculate eye-specific synapses in the mouse brain. The outcome of binocular synaptic competition was determined by the relative eye-specific maturation of presynaptic vesicle content. Genetic disruption of spontaneous retinal activity prevented subsynaptic vesicle pool maturation, recruitment of vesicles to the active zone, synaptic development and eye-specific competition. These results reveal an activity-dependent presynaptic basis for axonal refinement in the mammalian visual system.

**One-Sentence Summary:** Spontaneous activity regulates the nanoscale remodeling of presynaptic terminals underlying eye-specific synaptogenesis and competition.

## Main Text

The refinement of eye-specific projections to the dorsal lateral geniculate nucleus (dLGN) is a classic model system for investigating the role of spontaneous neural activity in synaptic competition during mammalian brain development (*1*–*4*). In the mouse, axonal refinement of eye-specific territories is a competitive process involving selective branch addition/elimination dependent on patterned spontaneous neural activity in the eyes (“retinal waves”) (*2*–*6*). Eye-specific refinement has been studied by imaging labeled axons, but direct analysis of synaptic competition has been hindered by difficulties in identifying immature retinogeniculate synapses based on ultrastructural features in electron microscopy (EM) images (*7*, *8*). Further, the diffraction limit of conventional light microscopy precludes fluorescence imaging analysis of eye-specific synaptic development. To address this gap, we used volumetric super-resolution microscopy in situ to measure nanoscale structural properties of retinogeniculate synapses during eye-specific competition in the mouse. We then compared synaptic refinement in wild-type (WT) mice with β2^-/-^ mice, a mutant strain with disrupted cholinergic retinal waves and eye-specific segregation defects caused by deletion of the β2 subunit of the nicotinic acetylcholine receptor (*5*, *9*).

To identify eye-specific synapses, we injected fluorescent anterograde tracer (cholera toxin subunit beta - Alexa Fluor 488, hereafter referred to as CTB) into the right eye and collected the left dLGN before (P2), during (P4), and toward the close (P8) of eye-specific segregation (Fig. 1A). We used a serial-section single-molecule localization imaging approach based on STochastic Optical Reconstruction Microscopy (STORM) to collect three-dimensional super-resolution fluorescence imaging volumes (~45K μm^3^ each) from the contralateral and ipsilateral regions of each dLGN sample (Fig. 1, A and B). For synapse identification, we immunostained tissue with antibodies against the presynaptic proteins vesicular glutamate transporter 2 (VGluT2) and Bassoon together with the postsynaptic protein Homer 1 (Fig. 1C). Eye-specific contralateral-eye synapses were CTB (+) while ipsilateral-eye synapses were CTB (-) (Fig. 1C). Binocular control injections revealed that 90% of dLGN VGluT2 clusters were CTB (+), demonstrating high efficiency synapse labeling during eye-specific segregation (P4-P8) (fig. S1A).

**Fig. 1.**
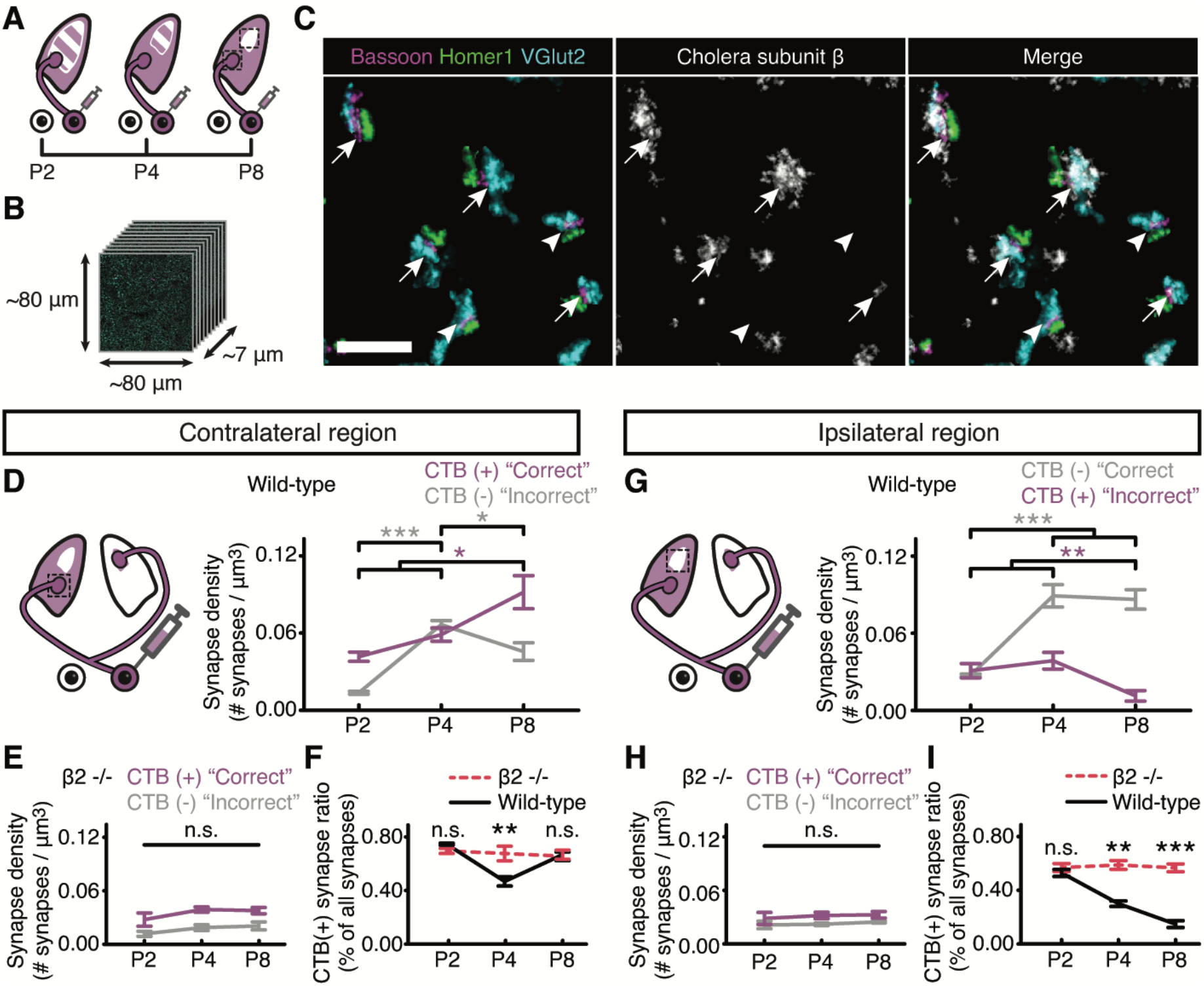
Spontaneous retinal activity regulates eye-specific synapse refinement. (**A**) Anterograde tracing identified eye-specific synapses in two regions of interest at three developmental time points. (**B**) STORM image stacks were ~45K μm^3^. (**C**) Representative STORM synapse (left), CTB signal (middle), and merged (right) images from P8 WT dLGN highlight CTB (+) contralateral-eye (arrows) and CTB (-) ipsilateral-eye synapses (arrowheads). Scale bar, 1 μm. (**D** to **I**) Synapse density in the contralateral (D to F) and ipsilateral (G to I) regions. CTB (-) ipsilateral synapse density increases (P2-P4) in both contralateral (D) and ipsilateral (G) regions. CTB (-) “incorrect” synapses (contralateral region, D) and CTB (+) “incorrect” synapses (ipsilateral region, G) are then lost (P4-P8). β2^-/-^ mice showed no differences in eye-specific synapse densities (E and H) and the ratio of CTB (+) synapses in each eye-specific region was constant (F and I). Error bars, mean ± SEM. N = 3 mice for each group. *P < 0.05, **P < 0.01, ***P < 0.001, one-way ANOVA with post-hoc Tukey test.

We quantified developmental changes in eye-specific synapse density within the contralateral region (Fig. 1, D to F), where the density of “correct” CTB (+) contralateral-eye synapses increased progressively and peaked at P8 (Fig. 1D). In the same region, the density of “incorrect” CTB (-) ipsilateral-eye synapses increased from P2 to P4 (Fig. 1, D and F), consistent with delayed ipsilateral axon ingrowth (*6*). The formation of ipsilateral-pathway synapses increased eye-specific synapse elimination resulting in a ~31% reduction in “incorrect” CTB (-) synapse density from P4-P8 (Fig. 1, D and F). Within the ipsilateral region (Fig. 1, G to I), the density of “correct” CTB (-) ipsilateral-eye synapses increased from P2-P4 and was stable from P4-P8 (Fig. 1G). Following ipsilateral synaptogenesis, “incorrect” CTB (+) contralateral-eye synapse density decreased ~72% between P4-P8 (Fig. 1, G and I). In contrast to WT, synapse development was disrupted in β2^-/-^ mice (Fig. 1, E and H) and the ratio of eye-specific synapses within each region did not change (Fig. 1, F and I). Cell body density and neuropil fraction were the same between wild-type and β2^-/-^ animals across all ages (fig. S1, B and C), suggesting normal dLGN expansion and a specific failure of synapse development in β2^-/-^ mice.

To test if reduced eye-specific synapse density resulted from defects in presynaptic or postsynaptic machinery necessary for neurotransmission, we quantified the size (volume) and protein enrichment (total signal intensity) of eye-specific VGluT2, Bassoon, and Homer1 clusters in the contralateral region (Fig. 2A-D; Fig. S2A-C). In WT and β2^-/-^ mice, presynaptic clusters (VGluT2 and Bassoon) grew larger (Fig. 2, B and C) and contained more proteins (fig. S2, A and B) over development but the effect size was greater for WT (Fig. 2B and fig. S2A). In contrast, postsynaptic Homer1 clusters in WT mice decreased in size from P2-P8 (Fig. 2D) with no change in protein enrichment (fig. S2C). Synaptogenesis in WT mice between P2-P4 (Fig. 1, D and G) generated nascent Homer1 clusters that were smaller and less enriched compared with β2^-/-^ mutants at comparable ages (Fig. 2D and fig. S2C).

**Fig. 2.**
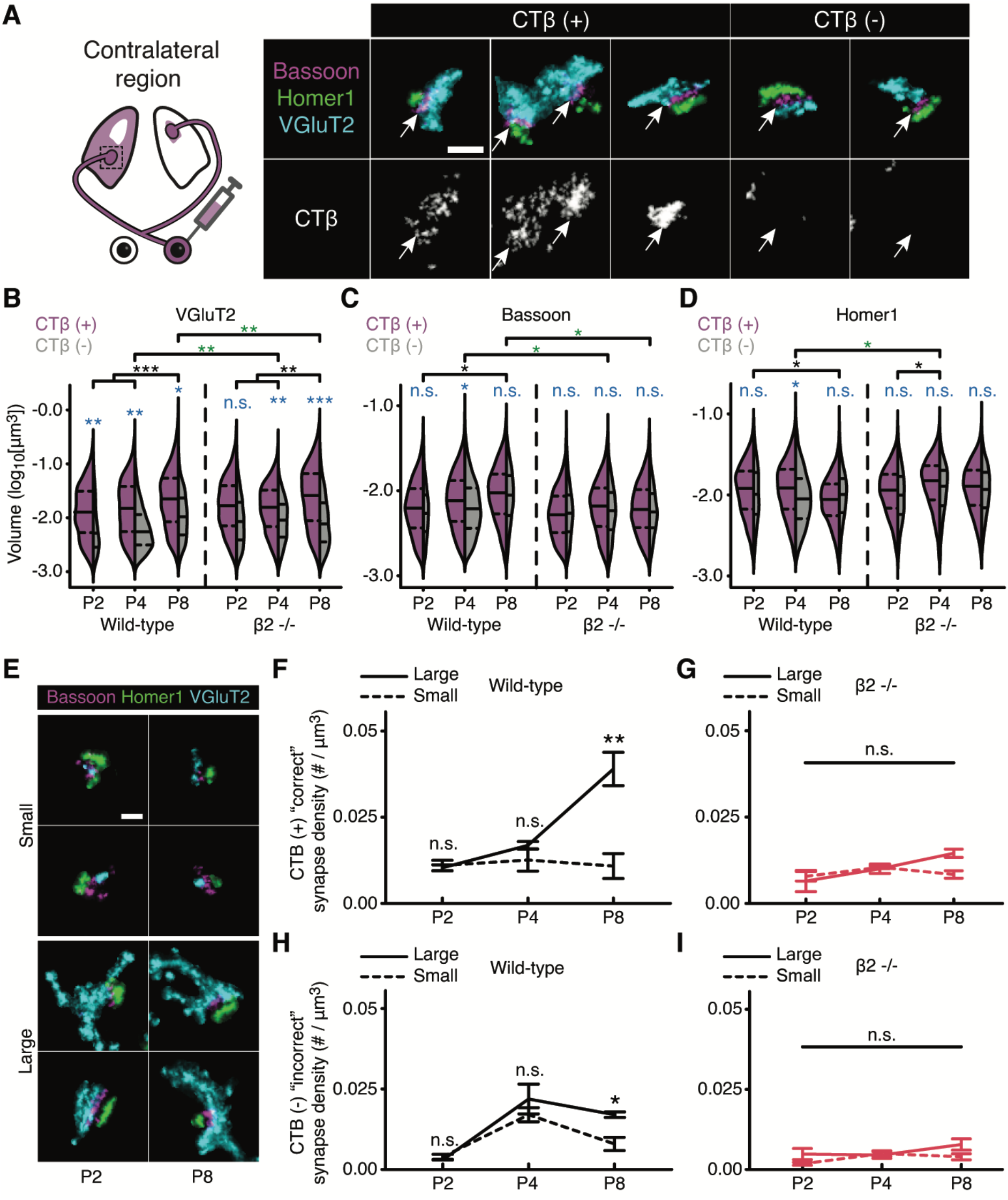
Presynaptic maturation underlies eye-specific competition. (**A**) CTB signals (arrows) define eye-specific synapses. (**B** to **D**) VGluT2 and Bassoon volumes both increase and Homer1 volume decreases in WT (P2-P8). VGluT2 volume, but not Bassoon or Homer1, increases in β2^-/-^ mice (P2-P8). Solid lines, medians. Dashed lines, 25% and 75% intervals of the distribution. Blue asterisks/text, comparison between CTB (+) and CTB (-) clusters. Black asterisks, comparison within genotype across ages. Green asterisks, comparison between genotypes at equivalent ages. (**E**) Representative small / large synapses from WT mice at P2 (left) and P8 (right). (**F** to **I**) In WT mice, large CTB (+) “correct” synapses increased with no change in small synapse density (F). Small / large CTB (-) “incorrect” synapse densities decreased from P4-P8 in WT mice and the reduction was greater for small synapses (H). β2^-/-^ mice showed no differences in small / large synapse densities (G and I). Error bars, mean ± SEM. N = 3 mice for each age/genotype. *P < 0.05, **P < 0.01, ***P < 0.001. (B to D) Mixed model ANOVA and post-hoc Bonferroni’s test. (F to H) One-way ANOVA with post-hoc Tukey test. (A and E) Scale bars, 300 nm.

We next compared synaptic cluster volume and protein enrichment for CTB (+) versus CTB (-) synapses across ages/genotypes in the contralateral region. In WT mice, “correct” CTB (+) contralateral-eye synapses contained more VGluT2 protein than “incorrect” CTB (-) ipsilateral-eye synapses (Fig. 2B and fig. S2A). Eye-specific VGluT2 differences were also seen in β2^-/-^ mutants but were of smaller magnitude (Fig. 2B and fig. S2A). In contrast, presynaptic Bassoon and postsynaptic Homer1 clusters showed eye-specific differences only at P4 in WT mice (Fig. 2, C and D, fig. S2B) consistent with the peak of ipsilateral synaptogenesis (Fig. 1, D and G).

Differences in presynaptic vesicle content between “correct” versus “incorrect” eye-specific synapses suggested that vesicle pool size could be a determinant for synaptic maintenance. To test this, we measured presynaptic VGluT2 volume distributions for all retinogeniculate synapses and found these were well-represented by a two-peak Gaussian fit for each age/genotype (fig. S3, A and B; R^2^ >0.96 for all fits). We defined VGLuT2 clusters with volumes smaller than the lower peak as a “small” synapse population and those larger than the upper peak as a “large” synapse population (fig. S3B). The synapse classification, defined by peak positions, was stable across ages/genotypes (fig. S3C) and large/small eye-specific VGluT2 clusters were morphologically similar across all conditions (Fig. 2E). The developmental increase in VgluT2 cluster volume for “correct” CTB (+) synapses (Fig. 2B) resulted from an increased density of large synapses while small synapse density remained stable (Fig. 2F). In contrast, ipsilateral-pathway synaptogenesis from P2-P4 increased the density of both large and small “incorrect” CTB (-) synapses (Fig. 2H). The density of both large and small CTB (-) synapses later decreased between P4-P8, but the decrease in small synapses (~50%) was more significant compared with larger synapses (~17%) (Fig. 2H) indicating “incorrect” synapses with greater presynaptic vesicular content were more likely to survive eye-specific competition. Developmental changes in the density of large versus small synapses of either eye of origin were not seen in β2^-/-^ mutants (Fig. 2, G and I), demonstrating an activity-dependent failure of presynaptic vesicle pool maturation underlying eye-specific synaptic refinement.

To further examine subsynaptic differences in vesicle organization and protein content underlying synaptic competition, we measured VGluT2 signal volume and enrichment within a 70 nm shell surrounding each active zone (AZ) Bassoon cluster to quantify putative docked vesicles (Fig. 3A). In WT mice, the volume and protein enrichment of docked VGluT2 signal increased over development for both CTB (+) and CTB (-) synapses in the contralateral eye-specific region (Fig. 3B and fig. S4A). No changes in docked VGluT2 were detected in β2^-/-^ mice across development (Fig. 3B and fig. S4A). To rule out the possibility that the measured increase in docked vesicle signal in WT animals was caused by the developmental increase in Bassoon cluster area and shell volume (Fig. 2C), we normalized docked VGluT2 signals to the total shell volume and again found a developmental increase in docked VGluT2, indicating that in WT, but not β2^-/-^ mice, vesicle proteins condensed near the presynaptic AZ (fig. S4, B and C). Interestingly, not all AZ Bassoon clusters were associated with docked VGluT2 signal (Fig. 3C). In WT mice prior to eye-specific synaptic refinement (P2), the number of VGluT2 ‘null’ synapses was higher within the “incorrect” CTB (-) synaptic population compared to “correct” CTB (+) synapses (Fig. 3D), demonstrating an early bias in eye-specific competition. In WT mice, the number of ‘null’ synapses from both eyes decreased over time while in β2^-/-^ mutants there were no differences in ‘null’ synapse numbers regardless of age or eye of origin, demonstrating an activity-dependent failure of vesicle recruitment to the AZ (Fig. 3D).

**Fig. 3.**
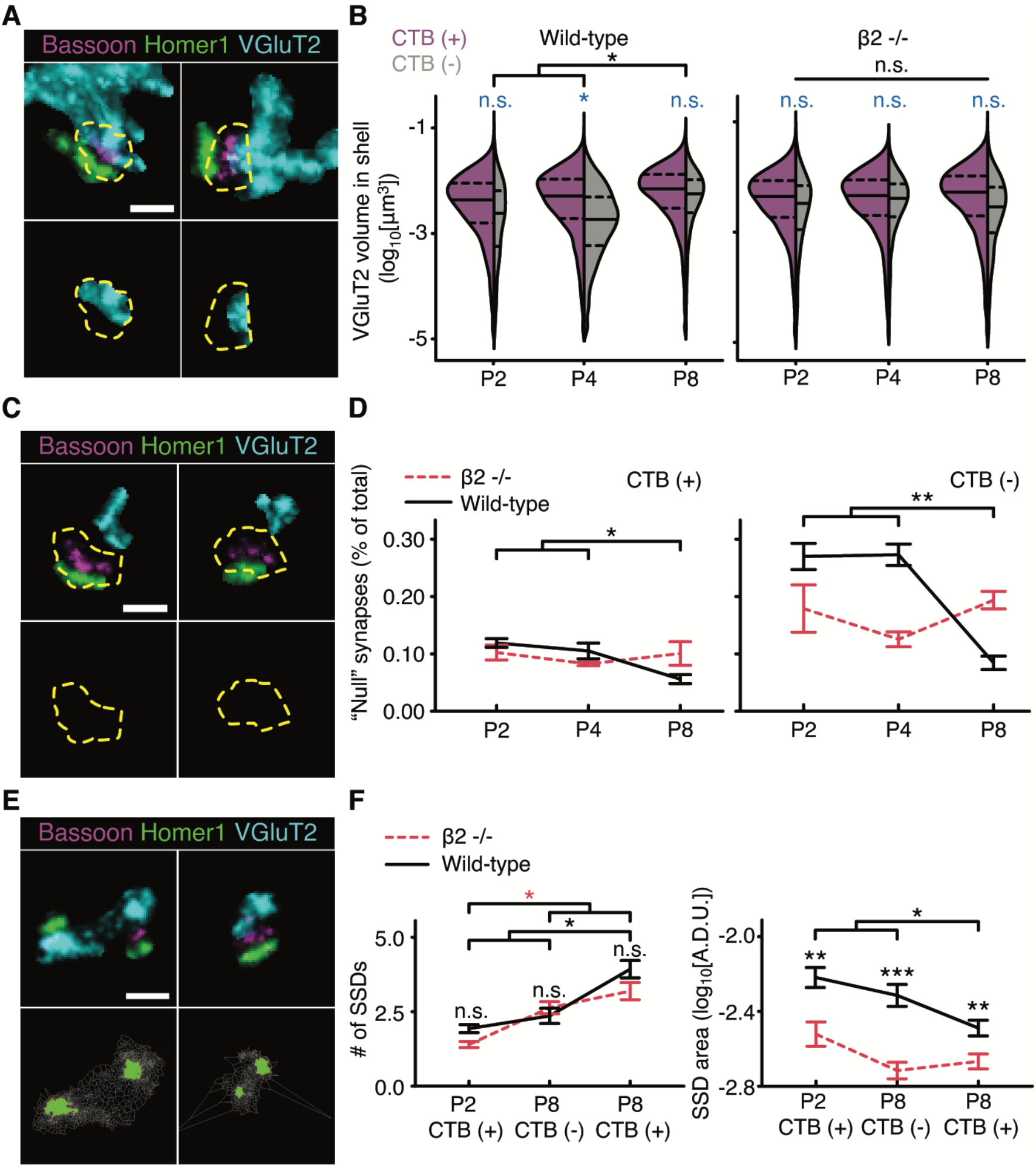
Activity-dependent maturation of subsynaptic vesicle organization during eye-specific competition. (**A**) VGluT2 signal was identified within a 70nm shell surrounding the AZ protein Bassoon. (**B**) In WT mice (left), docked VGluT2 volumes were largest at P8 while β2^-/-^ mice (right) showed no differences in docked vesicle volumes (right). Solid lines, medians. Dashed lines, 25% and 75% intervals of the distribution. (**C**) Representative ‘null’ synapses (**D**) In WT mice, the CTB (+) “null” synapse ratio decreased by P8 while β2^-/-^ animals showed no change (left). WT CTB (-) synapses contained a larger fraction of ‘null’ synapses that decreased by P8 in WT with no change in β2^-/-^ mice (right). (**E**) Representative VGluT2 SSDs identified by Vöronoi tessellation. (**F**) The average SSD number per VGluT2 cluster increased during development (left). In WT mice, but not in β2^-/-^ mice, the average SSD area decreased (right). In (D) and (F), error bars, mean ± SEM. *P < 0.05, **P < 0.01, ***P < 0.001. (B) Mixed model ANOVA and post-hoc Bonferroni’s test. (D) One-way ANOVA with post-hoc Tukey test. (F) Kruskal-Wallis test. (A, C, and E) Scale bars, 300nm.

To further investigate subsynaptic domains (SSDs) during eye-specific development, we used a Voronoï tessellation method (*10*) to measure SSD properties in eye-specific synapses within the contralateral region (Fig. 3E). In WT mice the average number of VGluT2 SSDs per synapse increased from P2-P8 (Fig. 3F), consistent with the increase in VGluT2 signal volume for both “correct” and “incorrect” eye-specific synapses (Fig. 2B). Concurrently, the average volume of individual VGluT2 SSDs decreased (Fig. 3F) while the measured number of localizations within individual VGluT2 SSDs did not change for synapses of either eye (fig. S5), suggesting that SSDs undergo spatial compaction during maturation. In β2^-/-^ mice, the average number of VGluT2 SSDs per synapse also increased, but compared with WT mice, SSDs were smaller across all ages (Fig. 3F). Consistent with our image analysis (Fig. 2), we detected no developmental maturation patterns or activity-dependent changes in either Bassoon or Homer1 SSDs during eye-specific segregation (fig. S5).

Eye-specific retinogeniculate refinement has been hypothesized to result from Hebbian synaptic plasticity where coactive synaptic outputs from neighboring retinal ganglion cells within each individual eye are strengthened, while uncorrelated outputs between the two eyes are weakened (*2*). For such a mechanism to work, the proper development of presynaptic release machinery is essential (*11*–*13*). Our data reveal that eye-specific presynaptic maturation depends on normal retinal wave activity as β2^-/-^ mice showed defects in synaptic refinement, reduced presynaptic vesicle content, and impairment of vesicle recruitment to the AZ. This is consistent with studies of activity-dependent synaptic competition at the neuromuscular junction where presynaptic disruption of neurotransmission leads to a competition bias between motor neuron axons followed by synapse loss (*14*). The activity-dependent maturation of vesicle pools underlying eye-specific competition is further consistent with electron microscopy results showing that “correct” eye-specific synaptic terminals contain more presynaptic vesicles versus “incorrect” terminals with no differences in PSD length/width (*7*).

The measured defects in eye-specific synaptogenesis and presynaptic terminal development support the synaptotropic hypothesis of neurite development (*15*) where synaptogenesis regulates the formation and stability of neurite branches (*16*, *17*). Under this model, the failure of presynaptic maturation and synaptic refinement in β2^-/-^ mice prevents axon branch stabilization and results in enlarged axon arbors (*5*). Defects in spontaneous retinal activity that impair presynaptic vesicle pool development would be expected to reduce neurotransmission efficacy, degrade postsynaptic coincidence detection, disrupt synaptogenesis, and expand individual axonal arbors resulting in a failure of eye-specific refinement based on Hebbian mechanisms.

These results explain axon segregation defects observed in previous imaging studies and highlight the emergence of super-resolution microscopy as a tool for investigating pathway-specific synaptogenesis in the developing brain.

## Acknowledgments

The authors thank Dr. Michael C. Crair for generously sharing the β2^-/-^ mouse line used in this work. We thank the members of the Speer lab for helpful discussions.

## Funding

National Institutes of Health grant DP2MH125812 (CMS)

Institutional Startup support provided by the University of Maryland

## Author contributions

Conceptualization: CZ and CMS

Data Curation: CZ and CMS

Formal Analysis: CZ and CMS

Funding Acquisition: CMS

Investigation: CZ and CMS

Methodology: CZ and CMS

Project Administration: CZ and CMS

Resources: CZ and CMS

Software: CZ and CMS

Supervision: CMS

Validation: CZ and CMS

Visualization: CZ and CMS

Writing – Original Draft Preparation: CZ and CMS

Writing – Review & Editing: CZ and CMS

## Data and materials availability

All data are available in the main text or supplementary materials. Raw STORM data sets are available upon request.

## Supplementary Materials

### Materials and Methods

#### Animals

Wild-type C57BL/6J mice used in this study were purchased from the Jackson Laboratory (Stock Number 000664). β2^-/-^ mice were a generous gift of Dr. Michael C. Crair (Yale School of Medicine). All experimental procedures were performed under an animal study protocol approved by the Institutional Animal Care and Use Committee (IACUC) at the University of Maryland. Neonatal male and female mice were used interchangeably for all experiments. Tissue from biological replicates (N=3 animals) was collected for each age (P2/P4/P8) from each genotype (WT and β2^-/-^) (18 animals total).

#### Eye injections

Intraocular eye injections were performed one day before tissue collection. Briefly, mice were anesthetized by inhalant isoflurane (~2-5% at a flow rate of 32 ml/min) for 2-3 minutes using a SomnoSuite low-flow anesthesia system (Kent Scientific). Sterile surgical spring scissors were used to gently part the eyelid to expose the corneoscleral junction. A small hole was made in the eye using a sterile 34-gauge needle and ~1μl of cholera toxin subunit B conjugated with Alexa Fluor 488 (CTB-488, ThermoFisher Scientific, Catalogue Number: C34775) diluted in 0.9% sterile saline was intravitreally pressure-injected into the right eye using a pulled-glass micropipette coupled to a Picospritzer (Parker Hannifin). For control experiments to test CTB labeling efficiency, binocular injections were performed using identical volumes in each eye.

#### dLGN tissue preparation

Animals were deeply anesthetized with ketamine/xylazine and transcardially perfused with 5-10 mls of 37°C 0.9% sterile saline followed by 10 mls of room temperature 4% EM Grade paraformaldehyde (PFA, Electron Microscopy Sciences) in 0.9% saline. Brains were embedded in 2.5% agarose and sectioned in the coronal plane at 100μm using a vibratome. Cut sections were further fixed in 4% PFA for 30 minutes at room temperature and then washed for 30-40 minutes in 1X PBS. The dorsal lateral geniculate nucleus (dLGN) was identified by the presence of CTB-488 signals using a fluorescence dissecting microscope. A circular tissue punch with a diameter of ~ 500μm containing the dLGN was microdissected from each section using a blunt-end needle. Two tissue sections from the central portion of the dLGN along the anterior-posterior access were selected for all biological replicates.

#### Immunohistochemistry

dLGN tissue punches were blocked in 10% normal donkey serum (Jackson ImmunoResearch, Catalogue Number: 017-000-121) with 0.3% Triton X-100 (Sigma-Aldrich Inc.) and 0.02% sodium azide (Sigma-Aldrich Inc.) diluted in 1X PBS for 2-3 hours at room temperature and then incubated in primary antibodies for ~72 hours at 4°C. Primary antibodies used were Rabbit anti-Homer1 (Synaptic Systems, Catalogue Number: 160003, 1:100) to label postsynaptic densities (PSDs), mouse anti-Bassoon (Abcam, Catalogue Number AB82958, 1:100) to label presynaptic active zones (AZs), and guinea pig anti-VGluT2 (Millipore, Catalogue Number AB251-I, 1:100) to label presynaptic vesicles. Following primary antibody incubation, tissues were washed in 1X PBS for 6 x 20 minutes at room temperature and incubated in secondary antibody solution overnight for ~36 hours at 4°C. The secondary antibodies used were donkey anti-rabbit IgG (Jackson ImmunoResearch, Catalogue Number 711-005-152, 1:100) conjugated with Dy749P1 (Dyomics, Catalogue Number 749P1-01) and Alexa Fluor 405 (ThermoFisher, Catalogue Number: A30000), donkey anti-mouse IgG (Jackson ImmunoResearch, Catalogue Number 715-005-150, 1:100) conjugated with Alexa Fluor 647 (ThermoFisher, Catalogue Number: A20006) and Alexa Fluor 405, and donkey anti-guinea pig IgG (Jackson ImmunoResearch, Catalogue Number 706-005-148, 1:100) conjugated with Cy3b (Cytiva, Catalogue Number: PA63101). Tissues were washed 6 x 20 minutes in 1X PBS at room temperature after secondary antibody incubation.

#### Post-fixation, dehydration, and embedding in epoxy resin

Tissue embedding was performed as previously described (*18*). Tissues were post-fixed with 3% PFA + 0.1% GA (Electron Microscopy Sciences) in PBS for 2 hours at room temperature and then washed in 1X PBS for 20 minutes. To plasticize the tissues for ultrasectioning, the tissues were first dehydrated in a graded dilution series of 100% ethanol (50%/70%/90%/100%/100% EtOH) for 15 minutes each at room temperature and then immersed in a series of epoxy resin/100% EtOH exchanges (Electron Microscopy Sciences) with increasing resin concentration (25% resin/75% ethanol; 50% resin/50% ethanol; 75% resin/25% ethanol; 100% resin; 100% resin) for 2 hours each. Tissues were transferred to BEEM capsules (Electron Microscopy Sciences) that were filled with 100% resin and polymerized for 16 hours at 70°C.

#### Ultrasectioning

Plasticized tissue sections were cut using a Leica UC7 ultramicrotome at 70 nm using a Histo Jumbo diamond knife (DiATOME). Chloroform vapor was used to reduce compression after cutting. For each sample, ~70-100 sections were collected on a coverslip coated with 0.5% gelatin and 0.05% chromium potassium (Sigma-Aldrich Inc.), dried at 60 degrees for 15 minutes, and protected from light prior to imaging.

#### Imaging chamber preparation

Coverslips were chemically etched in 10% sodium ethoxide for 5 minutes at room temperature to remove the epoxy resin and expose the dyes to the imaging buffer for optimal photoswitching. Coverslips were then rinsed with ethanol and dH2O. To create fiducial beads for flat-field and chromatic corrections, we mixed 715/755nm and 540/560nm, carboxylate-modified microspheres (Invitrogen, Catalogue Number F8799 and F8809, 1:8 ratio respectively) to create a high-density fiducial marker and then further diluted the mixture at 1:700 with Dulbecco’s PBS to create a low-density bead solution. Both high- and low-density bead solutions were spotted on the coverslip (~0.7 ul each) for flat-field and chromatic aberration correction respectively. Excess beads were rinsed away with dH2O for 1-2 minutes. The coverslip was attached to a glass slide with double-sided tape to form an imaging chamber. The chamber was filled with STORM imaging buffer (10% glucose, 17.5μM glucose oxidase, 708nM catalase, 10mM MEA, 10mM NaCl, and 200mM Tris) and sealed with epoxy.

#### Imaging setup

Imaging was performed using a custom single-molecule super-resolution imaging system. The microscope contained low (4x/10x air) and high (60x 1.4NA oil immersion) objectives mounted on a commercial frame (Nikon Ti-U) with back optics arranged for oblique incident angle illumination. We used continuous-wave lasers at 488nm (Coherent), 561nm (MPB), 647nm (MPB), and 750nm (MPB) to excite Alexa488, Cy3B, Alexa647, and Dy749P1 dyes respectively. A 405 nm cube laser (Coherent) was used to reactivate Dy749P1 and Alexa647 dye photoswitching. The microscope was fitted with a custom pentaband/pentanotch dichroic filter set and a motorized emission filter wheel. The microscope also contained an IR laser-based focus lock system to maintain optimal focus during automatic image acquisition. Images were collected on 640*640-pixel region of an sCMOS camera (ORCA-Flash4.0 V3, Hamamatsu Photonics) with a pixel size of ~155 nm.

#### Automated image acquisition

Fiducials and tissue sections on the coverslip were imaged using the low magnification objective (4X) to create a mosaic overview of the specimen. Beads/sections were then imaged at high-magnification (60X) to select regions of interest (ROIs) in the Cy3B channel. Before final image acquisition, laser intensities and the incident angle were adjusted to optimize photoswitching for STORM imaging and utilize the full dynamic range of the camera for conventional imaging. Low-density bead images were taken in 16 partially overlapping ROIs. 715/755nm beads were excited using 750 nm light and images were collected through Dy749P1 and Alexa 647 emission filters. 540/560nm beads were excited using a 488 nm laser and images were collected through Alexa 647, Cy3B, and Alexa 488 emission filters. These fiducial images were later used to generate a non-linear warping transform to correct chromatic aberration. Next, ROIs within each tissue section were imaged at conventional (diffraction-limited) resolution in all four-color channels sequentially.

Following conventional image acquisition, a partially overlapping series of 9 images were collected in the high-density bead field for all 4 channels (Dy749P1, Alexa 647, Cy3B, and Alexa 488). These images were later used to perform a flat-field image correction of non-uniform laser illumination across the ROIs. Another round of bead images was taken as described above in a different ROI of the low-density bead field. These images were later used to confirm the stability of chromatic offsets during imaging. All ROIs within physical sections were then imaged by STORM for Dy749P1 and Alexa 647 channels. Images were acquired using a custom progression of increasing 405nm laser intensity to control single-molecule switching. 8000 frames of Dy749P1 channel images were collected (60 Hz imaging) followed by 12000 frames of 647 channel images (100 Hz). In a second imaging pass, the same ROIs were imaged for Cy3B and Alexa 488 channels, each for 8000 frames (60 Hz).

For each physical section of the dLGN, we imaged the ipsilateral and contralateral fields separately. ROI selection was guided by CTB-488 signals and a fiducial knife cut made on the tissue at the dorsal edge of dLGN.

#### Image processing

Single-molecule localization was performed using a previously described DAOSTORM algorithm (*19*). Molecule lists were rendered as 8-bit images with 15.5nm pixel size where each molecule is plotted as an intensity distribution with an area reflecting its localization precision. Low-density fiducial images were used for chromatic aberration correction. We localized 715/755 beads in Dy749P1 and Alexa 647 channels, and 540/560 beads in Alexa 647, Cy3B, and Alexa 488 channels. A third-order polynomial transform map was generated by matching the positions of each bead in all channels to the Alexa 647 channel. The average residual error of bead matching was<15 nm for all channels. The transform maps were applied to both 4-color conventional and STORM images. Conventional images were upscaled to match the STORM image size and chromatic aberration was corrected for all images. The method to align serial sections was previously described (*18*). STORM images were first aligned to their corresponding conventional images by image correlation. To generate an aligned 3D image stack from serial sections, we normalized the intensity of all Alexa 488 images and used these normalized images to generate both rigid and elastic transformation matrices for all four-color channels of both STORM and conventional data. The final image stack was then rotated and cropped to exclude incompletely imaged edge areas. Images of the ipsilateral regions were further cropped according to CTB-488 signals to exclude contralateral areas.

#### Cell body filter

The aligned STORM images had non-specific labeling of cell bodies in Dy749P1 and Alexa 647 channels corresponding to Homer1 and Bassoon immunolabels. To limit synaptic cluster identification to the neuropil region we identified cell bodies based on their Dy749P1 signal and excluded these regions from further image processing. STORM images were convolved with a Gaussian function (σ=140nm) and then binarized using the lower threshold of a two-level Otsu threshold method. We located connected components in the thresholded images and generated a mask based on components larger than e^11^ voxels. Because cell body clusters were orders of magnitude larger than synaptic clusters, the cell body filter algorithm was robust to a range of size thresholds. The mask was applied to images of all channels to exclude cell body areas.

#### Eye-specific synapse identification and quantification

To correct for minor variance in image intensity across physical sections, we normalized the pixel intensity histogram of each section to the average histogram of all sections. Image histograms were rescaled to make full use of the 8-bit range. Using a two-level Otsu threshold method, the conventional images were thresholded into three classes: a low-intensity background, low-intensity signals above the background representing non-synaptic labeling, and high-intensity signals representing synaptic structures. The conventional images were binarized by the lower two-level Otsu threshold, generating a mask for STORM images to filter out background signals. STORM images were convolved with 77.5nm Gaussian function and thresholded in the same manner as the conventional images. Following thresholding, connected components were identified in three dimensions using MATLAB ‘conncomp’ function. A watershedding approach was applied to split large clusters that were improperly connected. To distinguish non-specific immunolabeling from true synaptic signals, we quantified two parameters for each cluster: cluster volume and cluster signal density calculated by the ratio of within-cluster pixels with positive signal intensity in the raw STORM images. Two separate populations were identified in 2D histograms plotted from these two parameters. We manually selected the population with higher volumes and signal densities representing synaptic structures. To test the robustness of the manual selection, we performed multiple repeated measurements of the same data and discovered a between-measurement variance of <1% (data not shown).

To identify paired pre-(Bassoon) and post-synaptic (Homer1) clusters, we first measured the centroid-centroid distance of each cluster in the Dy749P1 (Homer1) and Alexa 647 (Bassoon) channels to the closest cluster in the other channel. We next quantified the signal intensity of each opposing synaptic channel within a 140 nm shell surrounding each cluster. A 2D histogram was plotted based on the measured centroid-centroid distances and opposing channel signal densities of each cluster. Paired clusters with closely positioned centroids and high intensities of apposed channel signal were identified using the OPTICS algorithm. In total we identified 49,414 synapses from WT samples (3 samples each at P2/P4/P8, 9 total samples) and 33,478 synapses in β2^-/-^ mutants (3 samples each at P2/P4/P8, 9 total samples). Retinogeniculate synapses were identified by pairing Bassoon (Alexa 647) clusters with VGluT2 (Cy3B) clusters using the same method as pre/post-synaptic pairing. Synapses from the right eye were identified by pairing VGluT2 clusters with CTB (Alexa 488) clusters. The volume of each cluster reflected the total voxel volume of all connected voxels and the total signal intensity was a sum of voxel intensity within the volume of the connected voxels.

#### VGluT2 population analysis

To identify ‘small’ versus ‘large’ VGluT2 clusters in each sample, we used the MATLAB ‘histfit’ function to smooth the VGluT2 cluster volume histogram by fitting it to the kernel density distribution. The smoothed curve was then fit to the equation:

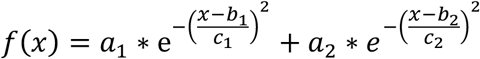

with the following boundary conditions:

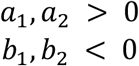

The peak positions were determined by the fitting results of b_1_ and b_2_.

#### Docked vesicle analysis

To identify putative docked vesicles based on VGluT2 signal, we computed a 70nm shell surrounding the surface voxels of each Bassoon cluster. VGluT2 signals within the shell area were reported as putative docked vesicle proteins. Bassoon clusters that did not contain VGluT2 signal withing the 70nm shell were defined as ‘null’ synapses.

#### Subsynaptic domain analysis

Based on our paired retinogeniculate eye-specific synapses, we analyzed the raw molecule localization distribution of each paired cluster using the SR-tessellation algorithm (*10*) for subsynaptic domain (SSD) analysis. We randomly selected 50 synapses each from P2 CTB (+), P8 CTB (-), and P8 CTB (+) synapse list for the analysis (N = 3 mice). The local density threshold was set to “2”, indicating that each selected SSD has a localization density 2x greater than the average density of the entire cluster. SSDs with fewer than 20 localizations were rejected. The area and number of localizations was quantified for each SSD identified in single 2D image planes through the center of each synapse.

#### Statistical analysis

Statistical analysis was performed using SPSS. Plots were generated by SPSS, R (ggplot2), or Python (Seaborn). We used a linear mixed model to compare Homer1, Bassoon, or VGluT2 cluster volumes and total signal intensity. In each comparison, the genotype, age, or eye of origin was a fixed main factor and biological replicate IDs were nested random factors. Comparison between main factor groups was performed by the post-hoc Bonferroni’s test. Cluster densities and SSD properties were compared by a one-way ANOVA test with Tukey’s post-hoc test (or Kruskal-Wallis test if the compared data sets did not pass the homogeneity of variance test). In violin plots, the solid line represents the median and dashed lines represent 25% and 75% intervals of the distribution. Stars in the figures indicate statistical significance: *=P<0.05, **=P<0.01, ***=P<0.001. Error bars represent standard error of the mean.

---

**Fig. S1.**
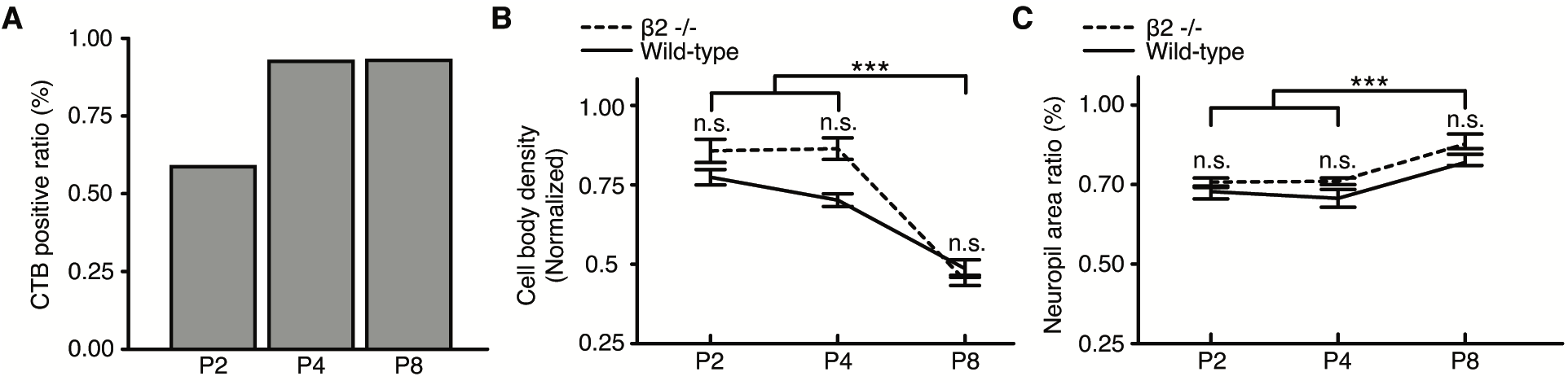
High efficiency eye-specific synapse labeling and homogeneous dLGN development for WT and β2^-/-^ mice during eye-specific segregation. (**A**) Colocalization of VGluT2 signal with CTB (+) labeling in binocular injection experiments was > 90% for P4 and P8 mice. (**B** and **C**) Cell body density measurements (B) and neuropil area ratios (C) were similar across development in WT and β2^-/-^ mice. Measurements reflect combined ipsilateral and contralateral regions. Error bars, mean ± SEM. N = 3 animals for each age/genotype. ***P < 0.001, one-way ANOVA with post-hoc Tukey test.

**Fig. S2.**
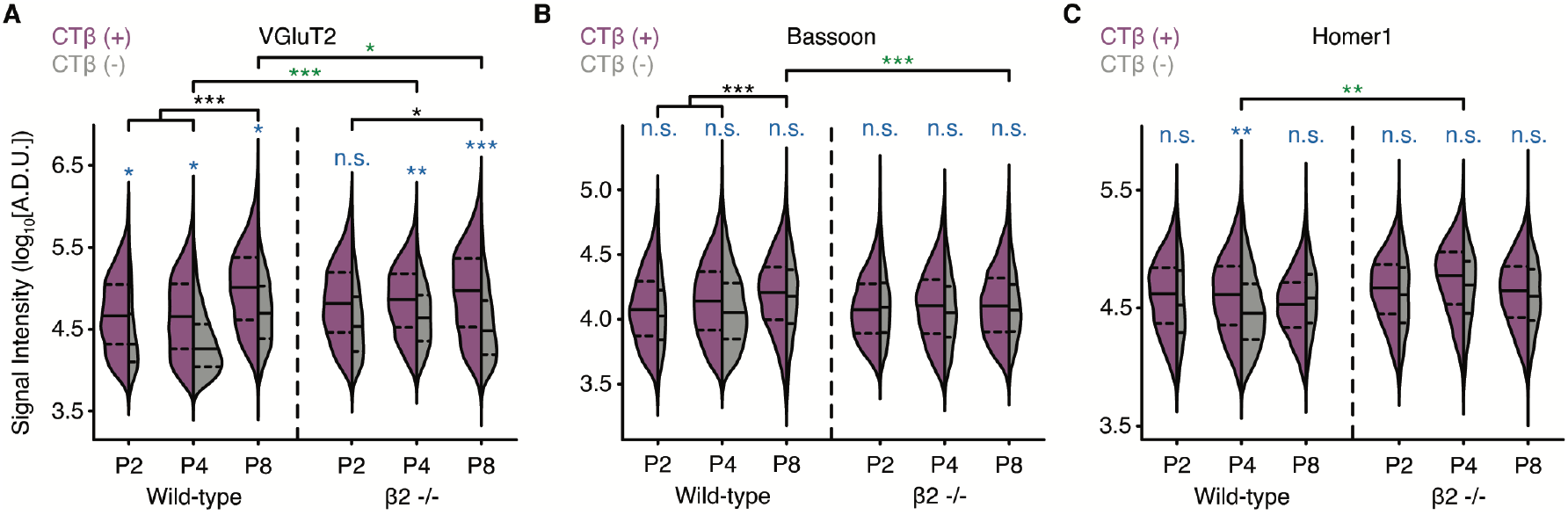
Developmental and activity-dependent increases in presynaptic signal intensity during eye-specific competition. (**A**) The mean VGluT2 cluster signal intensity increased over development in both WT and β2^-/-^ mice. Average WT VGluT2 signal intensity was lower at P4 but higher at P8 compared with β2^-/-^ mice. Eye-specific comparison revealed greater average signal intensity in CTB (+) “correct” clusters versus CTB (-) clusters across development for both genotypes except at P2 in β2^-/-^ mice. (**B**) The mean Bassoon cluster signal intensity increased over development in WT, but not β2^-/-^, mice. There were no differences in Bassoon cluster intensity distributions between CTB (+) “correct” and CTB (-) “incorrect” clusters for any age/genotype. (**C**) The mean Homer1 signal intensity did not change over development for either genotype. The mean Homer1 signal intensity was higher in β2^-/-^ mice at P4 than WT mice. Eye-specific mean signal intensity was greater in CTB (+) “correct” clusters versus CTB (-) “incorrect” clusters at P4 in WT, but not β2^-/-^, mice. Blue asterisks/text, comparison between CTB (+) and CTB (-) clusters. Black asterisks, comparison within genotype across ages. Green asterisks, comparison between genotypes at equivalent ages. Solid lines, medians. Dashed lines, 25% and 75% intervals of the distribution. *P < 0.05, **P < 0.01, ***P < 0.001, mixed model ANOVA with post-hoc Bonferroni’s test.

**Fig. S3.**
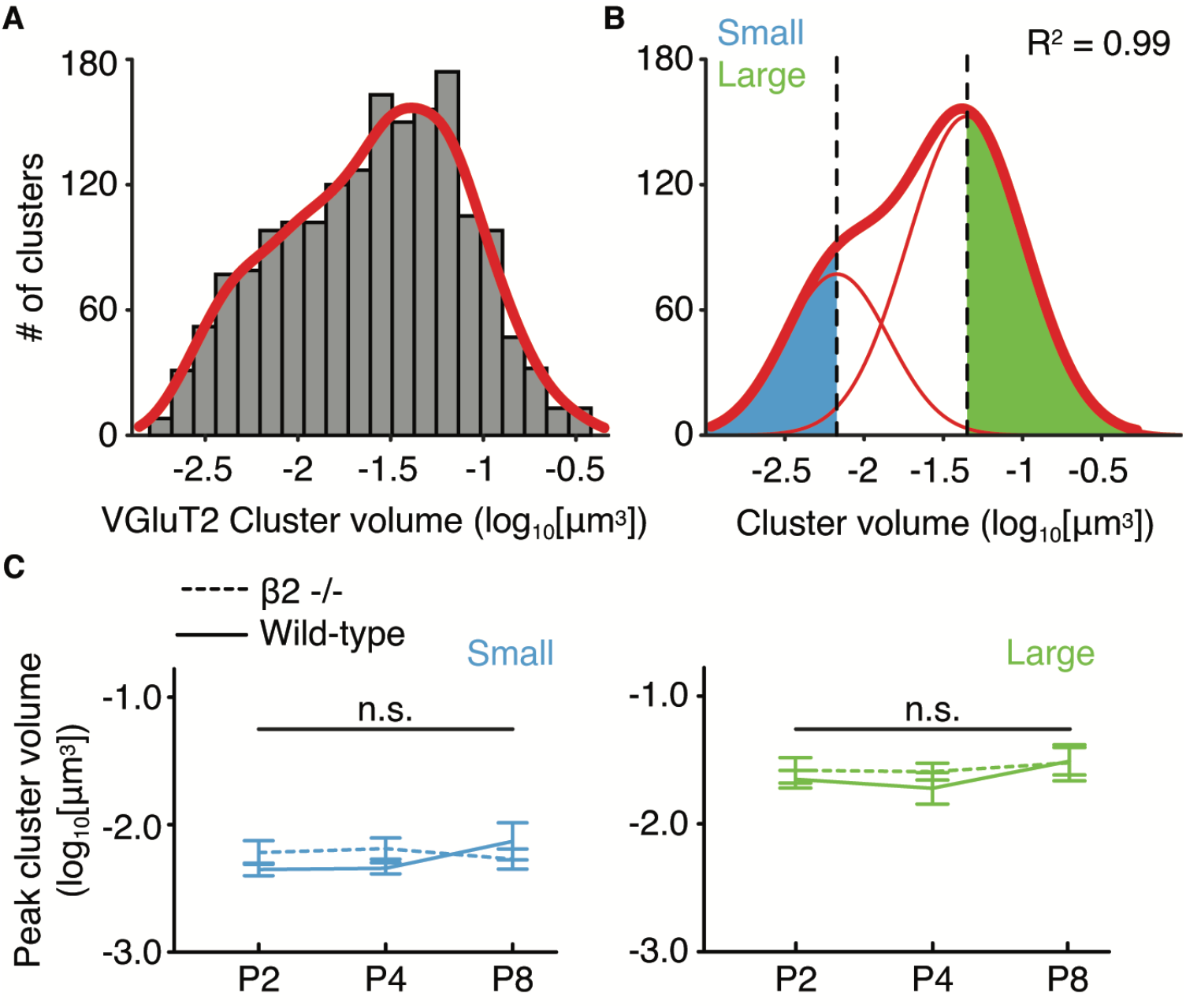
Two populations of presynaptic VGluT2 clusters in the mouse dLGN during eye-specific segregation. (**A**) Representative histogram of VGluT2 cluster volumes in the contralateral eye-specific region of a P8 WT mouse fit with the kernel density estimation (red line). (**B**) Definition of small and large VGluT2 clusters in the same sample as (A). The smoothed histogram (thick red line) was fit to 2-peak function (thin red lines) with peak positions marked by black dashed lines. Blue and green areas define small and large cluster volumes respectively. (**C**) Peak positions across all samples. The small population (left) and large population (right) peak positions were similar across all ages/genotypes. Error bars, mean ± SEM. One-way ANOVA test with post-hoc Tukey test.

**Fig. S4.**
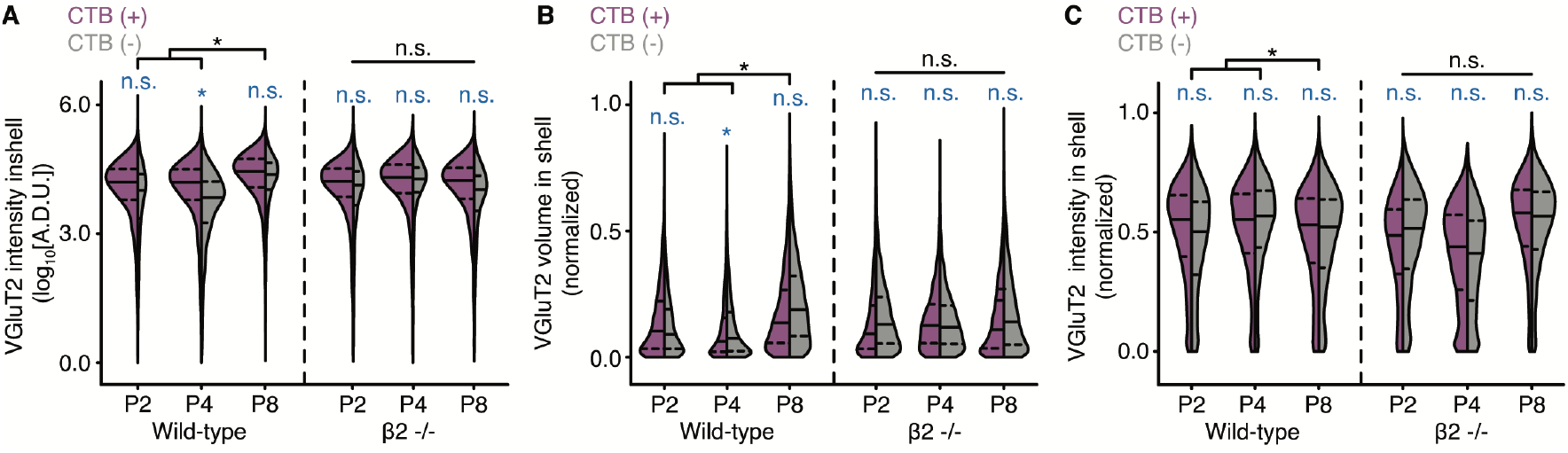
Activity-dependent maturation of docked vesicles during eye-specific competition. (**A**) Mean VGluT2 signal intensity within 70nm shell regions increased over development in WT (left), but not in β2^-/-^ (right), mice. (**B**) For each cluster, VGluT2 signal volume within the shell was normalized to the shell volume of the corresponding Bassoon cluster. Normalized VGluT2 shell volume increased over development in WT (left), but not in β2^-/-^ (right), mice. (**C**) For each cluster, VGluT2 intensity values within the shell were normalized to the shell volume of the corresponding Bassoon cluster. Normalized VGluT2 shell intensity increased over development in WT (left), but not in β2^-/-^ (right), mice. In all figures: solid lines, medians. Dashed lines, 25% and 75% intervals of the distribution. *P < 0.05. Mixed model ANOVA with post-hoc Bonferroni’s test.

**Fig. S5.**
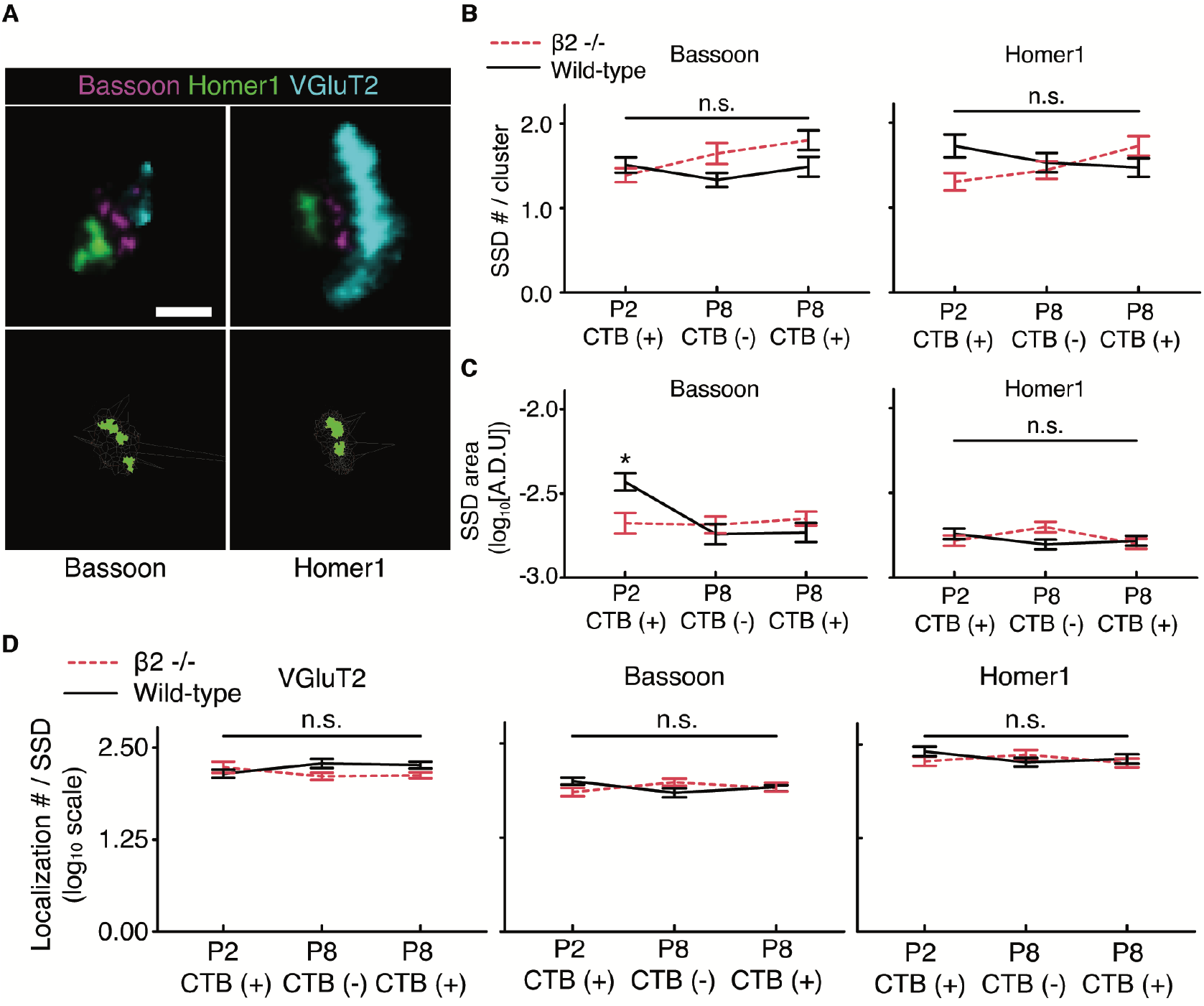
Active zone and postsynaptic protein SSDs were stable during eye-specific competition. (**A**) Representative SSDs within Bassoon (left) and Homer1 (right) clusters in the contralateral eye-specific region of a P8 WT mouse. (**B**) The number of Bassoon (left) and Homer1 (right) SSDs per synapse showed no developmental or activity-dependent changes across ages and genotypes. (**C**) SSD areas were similar across all ages and genotypes except that Bassoon SSDs were larger at P2 in WT compared to β2^-/-^ mice. (D) The average number of localizations per SSD in VGluT2 (left), Bassoon (middle), and Homer1 (right) clusters showed no developmental or activity-dependent changes. (B to D), error bars, mean ± SEM. *P < 0.05, Kruskal-Wallis test.

